# Topical Silver Diamine Fluoride for Dental Caries Arrest in Preschool Children: A Randomized Controlled Trial

**DOI:** 10.1101/131870

**Authors:** Peter Milgrom, Jeremy A. Horst, Sharity Ludwig, Marilynn Rothen, Benjamin W. Chaffee, Svetlana Lyalina, Katherine S. Pollard, Joseph L. DeRisi, Lloyd Mancl

## Abstract

**Objectives:** The Stopping Cavities Trial investigated effectiveness and safety of 38% silver diamine fluoride in arresting caries lesions.

**Materials and Methods:** Double-blind randomized placebo-controlled superiority trial with 2 parallel groups. Oregon <u>preschools.</u> 66 preschool children with ≥1 lesion. 38% silver diamine fluoride or placebo <u>(blue-tinted water),</u> applied topically to the lesion. The primary endpoint was caries arrest (lesion inactivity, Nyvad criteria) 14-<u>21</u> days post intervention. Dental plaque was collected from all children, and microbial composition was assessed by <u>RNA</u> sequencing from 2 lesions and 1 unaffected surface before treatment and at follow-up for 3 children from each group.

**Results and Conclusion:** Mean fraction of arrested caries lesions in the silver diamine fluoride group was higher (0.72; 95% CI; 0.55, 0.84) than in the placebo group (0.05; 95% CI; 0.00, 0.16). Confirmatory analysis using generalized estimating equation log-linear regression, accounting for the number of treated surfaces and length of follow-up, indicated the fraction of arrested caries was significantly higher in the treatment group (relative risk, 17.3; 95% CI: 4.3 to 69.4). No harms were observed. <u>RNA</u> sequencing analysis identified no consistent changes in relative abundance of caries-associated microbes, nor emergence of antibiotic or metal resistance gene expression. Topical 38% silver diamine fluoride was effective and safe in arresting cavities in preschool children. The treatment is applicable to primary care practice and may reduce the burden of untreated tooth decay in the population.

**Trial Registration:** ClinicalTrials.gov NCT02536040.

**Clinical Significance:** In this clinical trial, 72% of caries lesions were arrested by <u>silver diamine fluoride, with</u> no harms. <u>Contrary to the presumed antibacterial mechanism, lesion bacterial composition changed negligibly.</u> This simple topical treatment is applicable to primary care practice and may reduce the burden of untreated tooth decay in the population.

## INTRODUCTION

Dental and medical care providers are increasingly involved in early interventions to prevent tooth decay in young children.[1] The main preventive service is the topical application of fluoride varnish every 3 to 6 months. The rationale is that fluoride treatments plus anticipatory guidance may prevent the onset of tooth decay at the time of greatest risk. This is important because tooth decay negatively impacts <u>quality of life,[2,3]</u> particularly for children living in poverty, who have less access to dentists. Despite increased attention on young children, not all receive preventive treatments and large numbers still experience tooth decay. Primary care providers have had little option other than invasive and costly specialist treatment in the hospital under general anesthesia.

In this context, there is interest in simple treatments to halt progress of cavities (caries lesions) after tooth decay onset. Topical silver diamine fluoride is a clear liquid that is painted on the active lesion surface in milligram amounts and arrests the lesion. Cleared by the Food and Drug Administration in 2014 as a treatment for sensitive teeth and used off-label for the treatment of cavities in the United States since 2015, 12 clinical trials outside the United States have documented caries arrest<u>.[4-15]</u> Moreover, preventive benefits extend to unaffected teeth.[5,16] Serum concentrations of fluoride and silver after topical application revealed no potential toxicity.[17]

Silver ions are assumed to be primarily responsible for the antimicrobial action of silver diamine fluoride. Silver ions <u>can</u> lyse <u>all tested</u> bacteria, and denature enzymes that would breakdown collagenous dentin.[18,19] *Streptococcus mutans*, a primary pathogen in dental caries, is less able to form a biofilm on teeth treated *ex vivo*.[20] Fluoride promotes deposition of fluoroapatite, which is more resistant to acidic degradation than normal tooth structure.[21]

The purpose of this randomized controlled trial was to investigate the short-term effectiveness and safety of topically applied 38% silver diamine fluoride in arresting caries lesions in primary teeth.

## METHODS

### Design

Double-blind randomized placebo-controlled superiority trial with two parallel groups.

### Study Setting

<u>Three</u> Head Start programs <u>(preschools)</u> in <u>the U.S. state of</u> Oregon.

### Participants

The children were healthy by medical history; 24 to 72 months of age; and had at least one untreated cavitated active caries lesion with dentin exposed based on the Nyvad criteria <u>(level 3: “Enamel/dentin cavity easily visible with the naked eye; surface of cavity feels soft or leathery on gentle probing.”).</u>[22] The participant flow is outlined in Figure 1. Recruitment began February 25, 2016 and the final child was evaluated on May 10, 2016.

**Figure 1:**
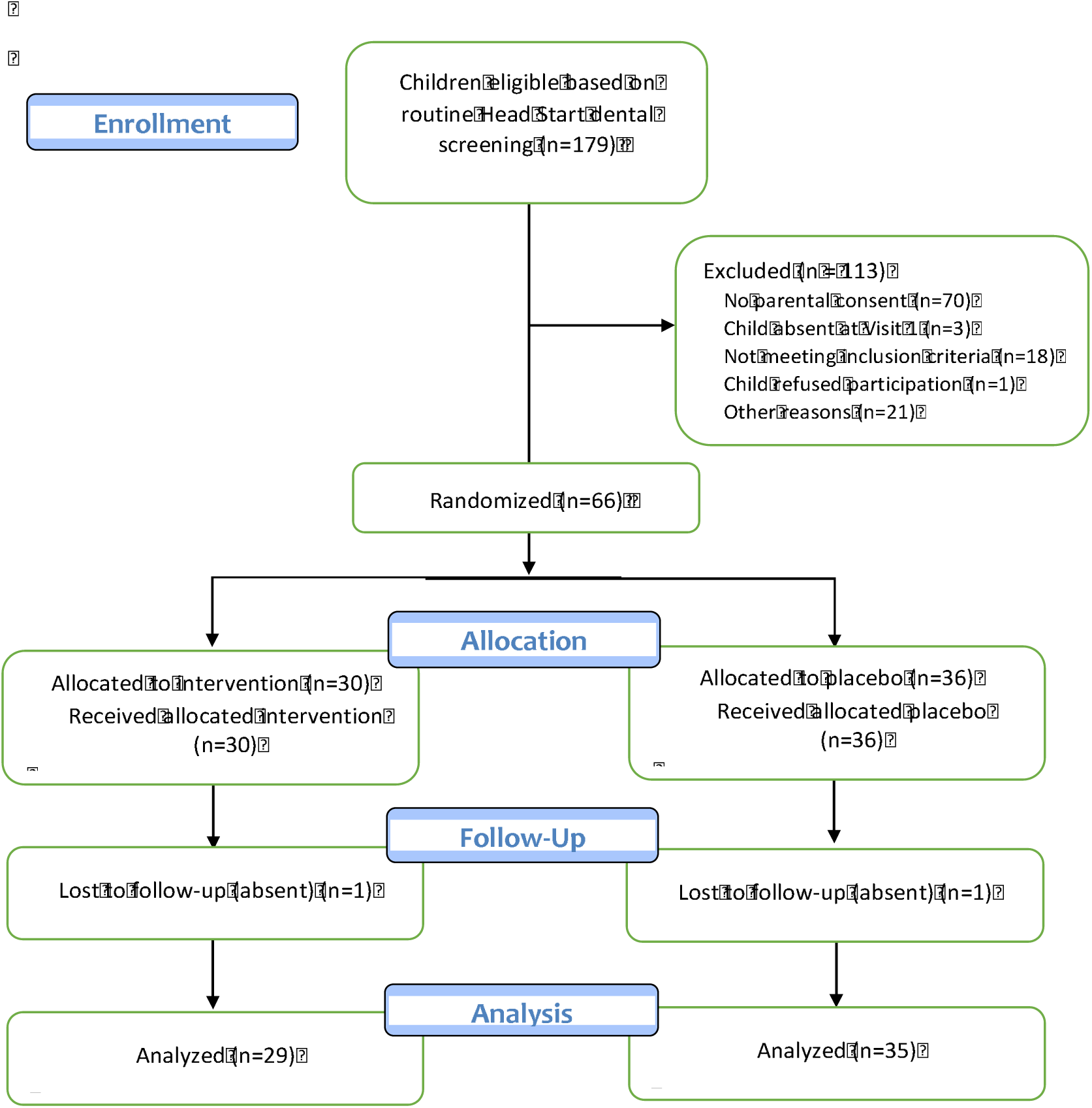
Flow chart of Stopping Cavities Trial

### Exclusion Criteria

Weight <15 kg; known sensitivity to silver or other heavy-metal ions; presence of any gingival or perioral ulceration or stomatitis; presence of a tooth abscess.

The study was conducted under IND 124808 from the Food and Drug Administration. The Western Institutional Review Board approved the protocol and the informed consent of participants was obtained. A $50 gift card was provided after follow-up.

### Treatment Conditions

Silver diamine fluoride (Advantage Arrest™, Elevate Oral Care LLC., West Palm Beach, FL). The test drug is a transparent blue-tinted solution containing 38 w/v% of silver (Ag) and 5.5 w/v% of fluorine (F). The placebo was non-fluoridated sterile water with a blue tint. The affected tooth surface was gently cleaned and dried with cotton gauze. The gingival tissue of the tooth was protected with petroleum jelly. An applicator was dipped into the agent and 3-4 mg applied to the lesion (1 drop treats 3 to 5 teeth). <u>No rinse or special instructions were performed following application.</u>

### Plaque Sample collection

Before treatment and at the beginning of the follow-up examination, plaque samples were taken from <u>up to</u> 2 caries lesions and one unaffected tooth surface, <u>selected sequentially from a list of tooth identifiers randomized for each patient,</u> by rubbing a 2-mm diameter plastic applicator on the lesion surface for 4 seconds. The applicator was then placed in 250μL tryptic soy broth with 15% glycerol and 2% sucrose, inverted twice, and kept at -20°C up to 2 months until moving to −80°C to await processing.

### Allocation

Participants were randomly assigned to groups with a 1:1 allocation using a computer-generated randomization schedule with stratification by <u>Head Start program</u> and randomly permuted blocks of sizes 2 and 4.

### Blinding

The dental providers who applied the treatments, participants, and examiners were blinded. Blinding was maintained by dispensing the test and comparator liquids from similar bottles. The test and placebo bottles were assigned codes A, B, C or D with 2 codes for test and 2 codes for placebo. <u>Four bottles were used to strengthen the blinding of treatment, and were otherwise identical.</u> The biostatistician knew only which two labels were the same, but was blinded to which labels were the test or placebo conditions.

### Primary Outcome

The primary outcome was caries lesion arrest. The hypothesis was that silver diamine fluoride is more effective than placebo in arresting caries lesions with dentin exposed in primary teeth. A visual/tactile examination was performed before and 14 to 21 days after treatment to identify and assess lesions for caries activity.[22] The teeth were examined with artificial light but without magnification in the preschool setting after drying with cotton gauze. A change from a Nyvad lesion code 3 to 6 <u>(“Enamel/dentin cavity easily visible with the naked eye; surface of cavity may be shiny and feels hard on probing with gentle pressure.”</u>[22]) was considered a positive outcome: caries arrest. The criteria have been shown to be reliable and valid.[22-25] The 14 to 21-day follow-up for outcome assessment was based on *in vitro* studies demonstrating silver diamine fluoride acts to arrest caries within this period.[26,27]

Six examiners were calibrated. The examiners reviewed the criteria and pictures showing the codes. Each independently examined 6 to 8 children with code 3 and code 6 lesions, including lesions treated with silver diamine fluoride. One investigator served as the index examiner. The observed agreement with the index examiner and 6 examiners ranged from 89.3% to 93.2% and the kappa statistic ranged from 0.68 to 0.79.

### Secondary outcome

The secondary outcome was to measure the change, if any, in relative abundance of microbial species or genus, assessed by metatranscriptomic (RNA) deep sequencing. The hypothesis was that caries-associated bacteria would decrease following treatment. We also hypothesized that prevalence of antibiotic and resistance genes would not change. RNA was used in proxy for vitality because many oral bacteria cannot be cultivated, and presence of RNA signifies production thereof by vital microbes within an hour beforehand. <u>Plaque samples were collected from all participants.</u> One participant in each study arm from each of the <u>3 study sites was selected from sequential lists of subject identification numbers by the biostatistician for the pilot analyses described here;</u> treatment arm participants were selected from the subset for whom all studied lesions successfully arrested <u>and placebo arm participants were selected from the subset from whom all studied lesions did not arrest.</u>

### Harms

Within 24 to 48 hours of treatment, trained staff contacted parent/caregivers by telephone about adverse events. The questions included (Figure S3): (1) Has your child required medical care since <u>his/her dental</u> visit <u>in the last 48 hours</u>? (2) Has your child been to an emergency room, medical doctor, nurse or health care provider? (3) Since receiving the solution on the teeth has your child experienced any of the following? Nausea; Not eating; Vomiting, Difficulty swallowing or breathing; Swelling around the lips or skin of the face; itchiness around the lips or skin of the face; hives or rash; stomach ache; or diarrhea.

At the follow-up visit 14 to 21 days after treatment, a dental provider performed a visual examination to detect gingival or soft tissue stomatitis or ulcerative lesions. Lesions were characterized as mild or severe and either <u>localized</u> or generalized.

### Analysis Plan

The primary outcome was the fraction of treated surfaces with arrested lesions at follow-up (code 6). The hypothesis was evaluated according to the intent-to-treat principle and based on all children who completed follow-up (97.0%). An exact two-sample permutation test was used to compare the mean fraction of arrested lesions between the two treatment conditions.[28] Bias-corrected bootstrap confidence intervals using 10,000 replications were also constructed for the mean fraction of arrested lesions by treatment condition and for the difference in mean fraction of arrested lesions between the 2 treatment conditions.[29,30] Log-linear regression, implemented using generalized estimating equation (GEE) methods, was used as a confirmatory analysis, comparing the rate of arrested lesions between the 2 treatment conditions, which accounted for the number of treated surfaces and length of follow-up by including the product of these 2 quantities as an offset term in the regression analysis.[31] In addition, the proportion of children with 100% of caries arrested was compared between the 2 treatment conditions using Fisher’s exact test, and exact 95% confidence intervals were constructed for the proportion by treatment condition and for the difference in proportions between the 2 treatment conditions.[32,33] A two-sided 5% significance level was used for all statistical inference. Statistical analyses were performed using SAS, Version 4 (SAS Institute Inc., Cary, NC, USA) and R, Version 3.3.0 (R Core Team, Vienna, Austria).

### Sample Size

Based on a two-sample t test, for 80% power it was estimated 79 to 100 participants per group were required to demonstrate a difference of 0.40 to 0.45 SDs in the mean fraction of arrested caries. If there were no arrested caries in the control group and 25% in the silver diamine fluoride group, it was estimated this difference would correspond to a difference of 0.40 to 0.45 SDs. Due to a delay in initiating the study and the need to complete follow-up by the end of the school year, only 66 participants were enrolled. However, the observed difference in the mean fraction of arrested caries was much higher (2.4 SDs) than used in the sample size determination.

### Metatranscriptomic sequencing library preparation

Total RNA was extracted by adding the applicator and 100 μL holding media to 400μL Trizol, bashing with 2mm ceramic beads for 3 minutes at 150 Hz with a TissueLyser (Qiagen, Hilden, Germany), and processing with an RNA Clean and Concentrator kit (Zymo Research, Irvine, CA, USA). Comparable amounts of RNA were obtained from each sample (1.5±1.1 μg per sample; mean ± standard deviation). Holding media, and that spiked with *Saccharomyces cerevisiae* RNA were used as controls.

5μg RNA was reverse transcribed to single-stranded complementary DNA (cDNA) using random hexamer primers and amplified to double-stranded (ds) cDNA (73±221 ng per sample) with New England Biolabs’ (NEB, Ipswitch, MA, USA) Ultra v1.5 Directional RNA Library Prep Kit,[34] then purified and size selected with 1.8x Ampure XP reagent (Beckman-Coulter Life Sciences, Indianapolis, IN, USA). The cDNA was converted to Illumina libraries using the NEBNext Ultra II DNA Library Prep Kit (E7645) according to the manufacturer’s recommendation with the addition of USER enzyme (NEB) when adding Illumina adapters, and purification steps with 0.9x Ampure. Dual index barcodes were added during 10 steps of PCR library enrichment. BioAnalyzer traces revealed high abundance 300-550 long oligonucleotides. Each sample was amplified again with dual-indexing primers on an Opticon qPCR machine (MJ Research, Waltham, MA, USA) using a Kapa Library Amplification Kit (Kapa Biosystems) until the exponential portion of the quantitative PCR signal was found, then cleaned with 0.8x Ampure. Samples were then quantified by qPCR (Kapa Biosystems, Wilmington, MA, USA) and pooled for each participant, then re-quantified with a ddPCR Library Quantification Kit (Bio-Rad, Hercules, CA, USA) and pooled again. Sequencing was performed on one lane in a HiSeq 4000 (Illumina, San Diego, CA, USA) using 135 bp paired-end sequencing.

### Sequencing read analysis

Following CASAVA quality filtering (Illumina, San Diego, CA, USA), human sequences (v38) were removed using the STAR alignment tool.[35] Low-quality sequences were removed with PriceSeq Filter.[36] Redundant reads were removed using CD-HIT-DUP.[37] Low-complexity sequences (Lempel-Ziv-Welch ratio of less than 0.42) were removed. Illumina adapter and X174 control sequences were removed using BowTie2.[38] A random subset of 300,000 reads was searched against a repeat-masked[39] subversion of the NCBI nonredundant nucleotide (nt) database (version July 2015) using GSNAPL,[40] and processed with internal scripts. The subset was searched against the BacMet database[41] using RapSearch2[42] to assess levels of metal- and antibiotic-resistance genes.

High quality sequencing reads were obtained in abundance (2.9±1.9*10^6 reads) for all but one sample (56,835 from the unaffected surface of participant 3058 at follow-up). 21.7±5.0% of filtered reads mapped to nt, which is typical for metatranscriptomic sequencing. The media negative control showed low background (937 reads), verifying the clinical source of reads for the other samples. The spiked positive control resulted in nearly all reads matching *S. cerevisiae* (89%) or fungal yeasts (3%), as expected.

### Microbiology

Taxonomic and resistance gene counts were analyzed for significant changes using LME4 (R Core Team). Nested grouping was used to represent the experimental design of paired before-after samples and paired lesion-plaque samples from the same patients. The regression design was: ~treatment+time+treatment:time, where the ß coefficient for treatment:time was assessed for changes. <u>False discovery rates were considered significant when the *P* value divided by the number of assessed taxa or genes (Bonferroni correction) were less than 0.01.</u>

## RESULTS

Sixty-six children were randomized. Thirty-six participants were randomly assigned to placebo, treated with liquid A or D, and 30 were assigned to silver diamine fluoride, treated with liquid B or C. Follow-up was completed on 64 (97%). One child in each group did not return for follow-up. In addition, one child in the placebo group was uncooperative at the follow-up examination and only 11 of 20 teeth, which included all 4 treated teeth, were examined. Participant characteristics are in Table 1.

**Table 1.**
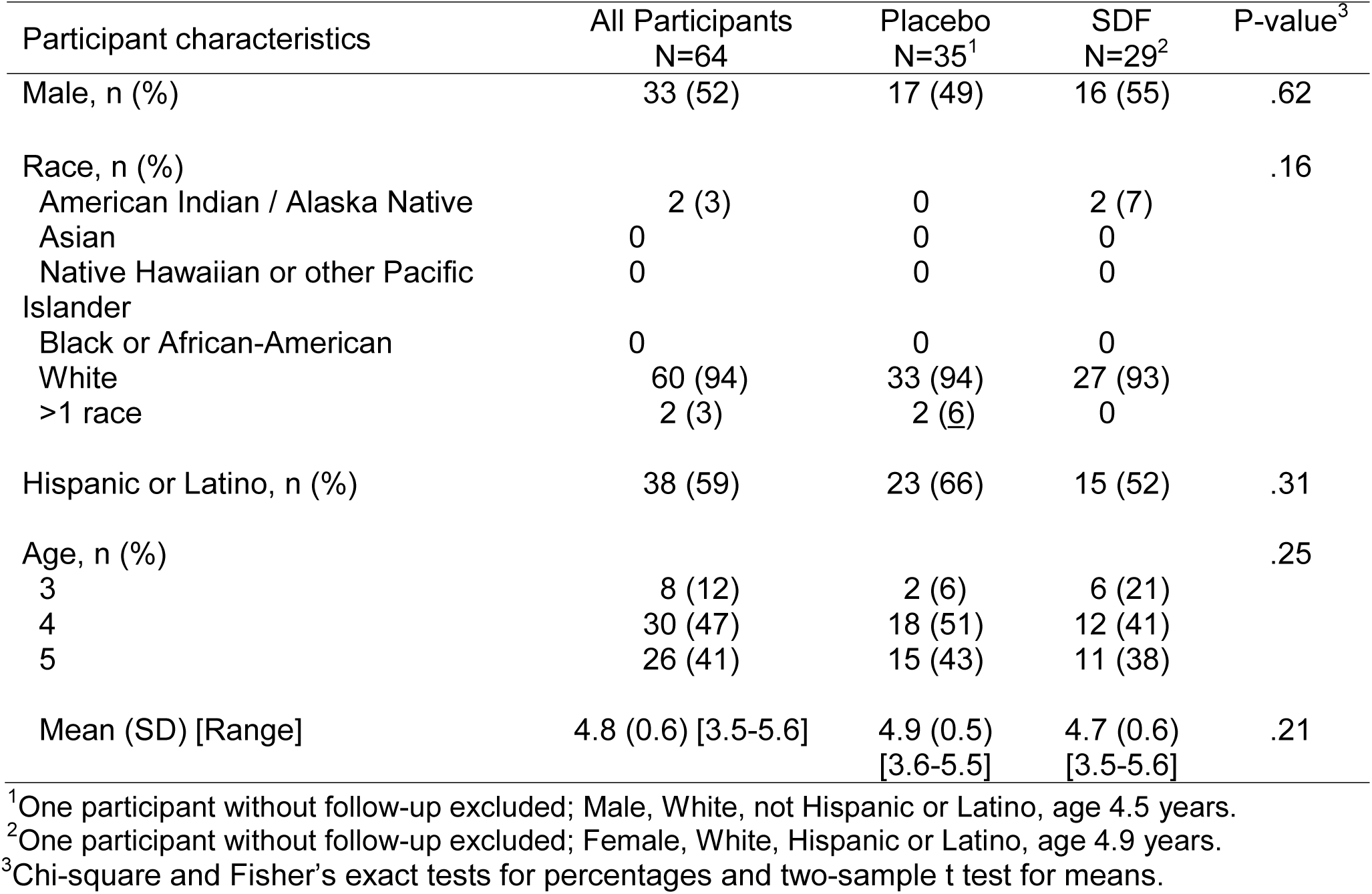
Participant characteristics in the Stopping Cavities trial of silver diamine fluoride (SDF) for the arrest of dental caries. <u>A subset of White children are also Hispanic or Latino, which is considered an ethnicity by the National Institutes of Health, rather than a separate race.</u>

Among the 64 children with follow-up, the mean (SD) number of treated teeth was 2.63 (1.09) in placebo and 2.83 (1.26) in silver diamine fluoride groups, respectively. The mean (SD) number of treated surfaces was 3.97 (2.56) in the placebo and 4.31 (2.74) in silver diamine fluoride groups, respectively. The mean (SD) length of follow-up was 17.2 (6.5) days in the placebo group and 16.4 (5.5) days in the silver diamine fluoride group (Table 2).

**Table 2.** Number treated teeth and surfaces and length of follow-up by treatment condition within the Stopping Cavities trial of silver diamine fluoride (SDF).

|  | N | Mean (SD) | Median (IQR) | Min - Max |
| --- | --- | --- | --- | --- |
| *Number of treated teeth* |  |  |  |  |
| Placebo | 35 | 2.63 (1.09) | 2 (2, 4) | 1 – 4 |
| SDF | 29 | 2.83 (1.26) | 3 (2, 4) | 1 – 4 |
| *Number of treated surfaces* |  |  |  |  |
| Placebo | 35 | 3.97 (2.56) | 4 (2, 4) | 1 – 14 |
| SDF | 29 | 4.31 (2.74) | 4 (2, 6) | 1 – 12 |
| *Length of follow-up, days* |  |  |  |  |
| Placebo | 35 | 17.2 (6.5) | 14 (14, 20) | 12 – 40 |
| SDF | 29 | 16.4 (5.5) | 14 (14, 14) | 12 – 28 |

### Outcomes

The mean fraction of <u>teeth with all lesions</u> arrested in the silver diamine fluoride group was 0.72 (95% CI; 0.55 to 0.84) and 0.05 in the placebo (95% CI; 0.00 to 0.16). The difference in mean fraction of <u>teeth with all lesions</u> arrested was 0.67; 95% CI: 0.49 to 0.80 (Table 3). GEE confirmatory analysis indicated the rate of arrest was significantly higher in the treated than in the placebo group (relative risk 17.3; 95% CI: 4.3 to 69.4).

**Table 3.** Fraction of arrested caries lesions by treatment condition in the Stopping Cavities trial with silver diamine fluoride (SDF).

| Condition | N | Mean (SD) | 95% CI<sup>1</sup> | Median (IQR) | Min - Max |
| --- | --- | --- | --- | --- | --- |
| SDF | 29 | 0.72 (0.38) | 0.55 to 0.84 | 1.00 (0.50, 1.00) | 0.0 – 1.0 |
| Placebo | 35 | 0.05 (0.18) | 0.00 to 0.16 | 0.00 (0.00, 0.00) | 0.0 – 1.0 |
| Difference<br>(Group 2 – Group 1) |  | 0.67 | 0.49 to 0.80 |  |  |

<sup>1</sup>95% CI: Bias-corrected bootstrap, 95% confidence interval for mean fraction of arrested caries and difference in mean fraction of arrested caries

Over half of children in the silver diamine fluoride group had 100% of lesions arrested (15/29; 51.7%; exact 95% CI: 32.5 to 70.6%) versus only 1 of 35 (2.9%; exact 95% CI: 0.00 to 14.9%) in the placebo group. The rate difference was 48.9%; exact 95% CI: 29.6 to 67.0%). (data not shown). No participant was identified with gingival or soft tissue stomatitis or ulcerative lesions.

### Microbiology

Metagenomic sequencing analysis revealed an abundance of vital microbes in all lesions, as inferred by presence of RNA. Contrary to the initial hypothesis, the relative abundance of only a few species were significantly altered in caries lesions by the intervention (Figure 2). No bacteria associated with caries changed significantly. *Streptococcus mutans* and *Lactobacilli*, the bacteria primarily associated with dental caries, were found in all samples except one unaffected surface before treatment, with no correspondence to treatment. All changes in relative abundance of species with a Bonferroni-adjusted probability of false discovery <0.01 were modest increases, not significant decreases as originally hypothesized. Low-level changes in many taxa were observed, but few were consistent. No significant changes in expression of antibiotic or metal resistance genes were observed.

**Figure 2.**
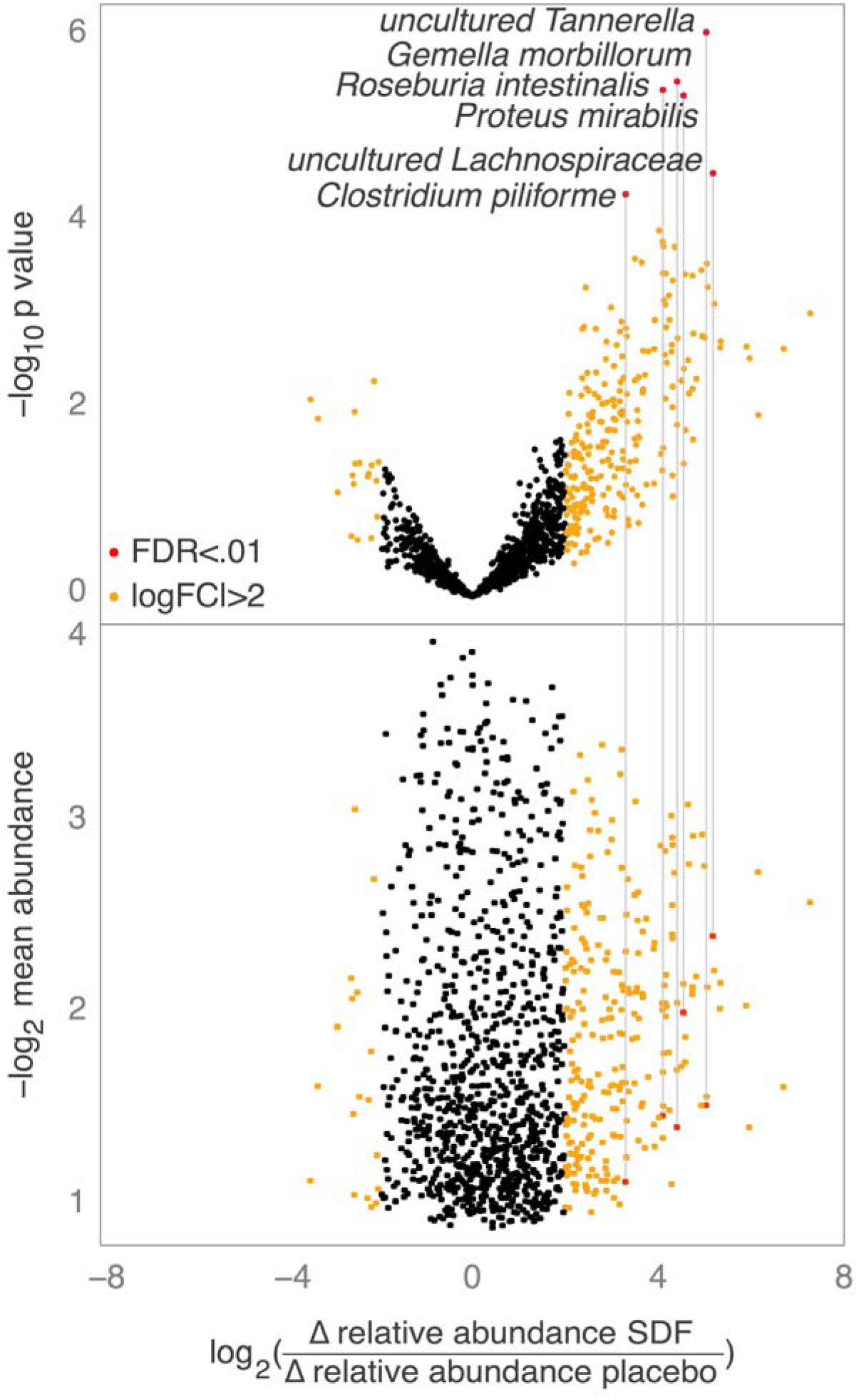
Changes in microbes following SDF treatment. Species are plotted on the horizontal axis by the log2 change in relative abundance following treatment in the SDF treatment group, relative to the placebo group. The top panel plots the p-value of the change, with labels for species with an estimated false discovery rate <0.01 after adjusting for multiple testing. The lower panel plots the mean abundance for each species. FDR: false discovery rate; FC: fold change; Δ: change; SDF: silver diamine fluoride.

Diversity analyses showed no significant differences (Supplemental Figure S1). Principal component analysis resulted in a highly-distributed explanation model (low attribution to any component), with some non-clustered outgrouping of samples in the silver diamine fluoride group after treatment (Supplemental Figure S2).

### Harms

Within 48 hours of treatment, staff successfully contacted 51 of the 66 (77%) parents by telephone. (Supplemental Table S1) The contact rates were 62.2% for the placebo group and 83.3% for the silver diamine fluoride group. Eight adverse events were reported by parents, 4 in each condition. Based on enrolled <u>participants,</u> the adverse event rate was 11.1% in the placebo group (exact 95% CI: 3.2 to 26.7%) and 13.3% in the silver diamine fluoride group (exact 95% CI: 3.7 to 30.7%). Based on all contacted <u>participants,</u> the adverse event rate was 15.4% in the placebo group (exact 95% CI: 4.3 to 34.9%) and 16.0% in silver diamine fluoride group (exact 95% CI: 4.5 to 36.1%). There were no statistically significant differences by treatment.

The type of adverse event, severity and relationship of the adverse event to the treatment condition are reported in Supplemental Table S2. The majority of the adverse events were either diarrhea or stomach ache, mild in severity and all resolved within 2 days of reporting.

## DISCUSSION

In 3 to 5-year-old children, 38% silver diamine fluoride was dramatically more effective than placebo clinically in arresting lesions after about two weeks and was safe. This study documents short-term effectiveness. Such treatments, if available from primary care providers, may prevent morbidity associated with untreated Early Childhood Caries and reduce treatment needs for patients awaiting specialist care in the hospital, or be a critical component of nonsurgical management.

Now with 9 published clinical trials in children from 18 to 36 months in duration, [4-12] many factors in optimizing the clinical protocol and regimen have been worked out: mechanical excavation of lesions with hand instruments or a dental drill is unhelpful,[43] a dose response is achieved by higher concentration (38% versus 12%),[7,9] and longer term effectiveness is greater if the treatment is repeated.[6,7,9] We also know that application only to lesions prevents lesions on untreated surfaces remarkably well,[5,43] and application to high risk surfaces that are not cavitated, such as the grooves of the biting surfaces in permanent molars, prevents caries directly.[11, 12] <u>Applying to all exposed surfaces, as is done with fluoride varnish, does not appear to be indicated.</u> No study has established the optimal retreatment interval; although once per 6 months achieves the highest efficacy.[5-7]

### Microbiology

Contrary to expectations, intensive microbiological analyses on a pilot set of patients showed no changes in the relative abundance of bacteria associated with dental caries (*P*>.05). This suggests that the microbial mechanism of caries arrest by silver diamine fluoride is indiscriminate and highly localized killing or inhibition of all bacteria in a caries lesion, rather than selection against any particular cariogenic bacteria. A leveling of bacterial groups is supported by trends in diversity analyses, with some level of change suggested by outgrouping of half the treated lesions in principal component analysis. This observation of increased bacterial diversity following treatment may be confirmed by analyzing more samples.

Dramatic decreases in microbial diversity following the use of traditional antibiotics enables the dangerous growth of opportunistic pathogens such as *Clostridium dificile.* The lack of any significant loss of bacterial species diversity here implies safety. Safety is further reassured by the lack of significant increases in disease-associated microbes, except *Proteus mirabilis*, a bacterium associated with urinary tract infections, which increased 17.2±1.7 fold (standard error SE; p=0.0013). Bacteria that may be protective against caries, such as *Streptococcus mitis*, opportunistically cause life-threatening infections. Our results strongly suggest that silver diamine fluoride treatment poses minimal risk of the unintended systemic effects that too often occur with other antimicrobials. These findings should be validated in more patients and with different sampling techniques.

### Harms

Adverse events occurring in the two days following application were similar between groups. One child in the treatment group was described as having a non-sore, non-irritated spot at the corner of the mouth that “looked like a burn,” which is a typical description of the silver precipitation that occurs upon contact with the skin, which resolved by the 2-week follow-up. <u>Discoloration was not considered a harm, as the Food and Drug Administration defines harm as “damage to health.”</u> No tooth pain was reported in any treatment group patients, despite deep lesions in some. No gingival or mucosal irritation was observed at follow-up in any patients.

### Limitations

The caries arrest outcome in this study was evaluated much earlier than in other clinical trials of silver diamine fluoride to treat dental caries; the shortest follow-up in other studies is 6 months. The average caries arrest rate in 6 months in these studies is roughly 43%.[5,6,8] The comparatively high rate observed in this study may be because the blue coloring of the agent allows more complete application or because there is a greater initial response that fades with time without reapplication. A limitation of the microbiological analyses is that they measure the relative abundance of various species rather than absolute concentrations within a lesion; previous observations suggest the quantity of microbes is dramatically reduced. Reactivation of lesions was observed in a 2-year study with only a single application.[9] Indeed, higher rates of caries arrest at 6 months were observed when silver diamine fluoride was applied thrice in two weeks versus once.[8] Monitoring of treated lesions to ensure arrest is indicated.

## CONCLUSION

Consistent with the literature, topical 38% silver diamine fluoride arrests tooth decay and is effective for the short-term treatment of dental caries in pre-school age children. The effect is rapid and safe. The potential for microbial resistance appears low.

## Acknowledgments

Ashley Danielson, Mindy Bentley, Stephanie Brouse, Kim Holliday, Tiffany Foy, and Brenna Southwick, administered treatments and served as examiners in the trial. Supported, in part, by NIDCR grant T32-DE007306 and funds from Advantage Dental Services, <u>LLC, and the NIH National Center for Advancing Translational Sciences through the University of Washington’s Clinical and Translational Science Award (CTSA), number UL1 TR000423.</u> Thanks to the DeRisi lab members, particularly Lillian Khan, Charles Langelier, Michael Wilson, Wei Gu, and Eric Chow for support in developing the sequencing protocol.

## Supplemental Tables

**Table S1.** Completion of 24-48 follow-up and reported adverse events.

|  | All<br>participants<br>N=66 | Group 1<br>N=35 | Group 2<br>N = 30 |
| --- | --- | --- | --- |
| 24-48 hour follow-up, n(%) |  |  |  |
| Completed | 51 (77.3) | 26 (62.2) | 25 (83.3) |
| Unable to contact | 15 (22.7) | 10 (27.8) | 5 (16.7) |
| Adverse event, n (%) | 8 (12.1) | 4 (11.1) | 4 (13.3) |

**Table S2.** Harms

| Study<br>identification | Group | Adverse event description | Severity<br>of event | Relationship of event<br>to study product |
| --- | --- | --- | --- | --- |
| 1015 | 1 | Diarrhea | Mild | Unrelated |
| 2004 | 1 | Diarrhea | Mild | Possibly related |
| 2082 | 1 | Stomach ache | Moderate | Possibly related |
| 3052 | 1 | Sporadic sore & hurting tooth;<br>Diarrhea | Mild | Probably not related |
| 1003 | 2 | Flu like symptoms (nausea, vomiting,<br>stomach ache, diarrhea) | Mild | Probably not related |
| 1007 | 2 | Redness around mouth, but not sore<br>or irritated | Mild | Probably not related |
| 1008 | 2 | Spot on corner of lip; looked like a<br>burn but was flat, not sore or irritated | Mild | Possibly related |
| 3056 | 2 | Nausea and stomach ache | Mild | Probably not related |

## Supplemental Figures

**Figure S1.**
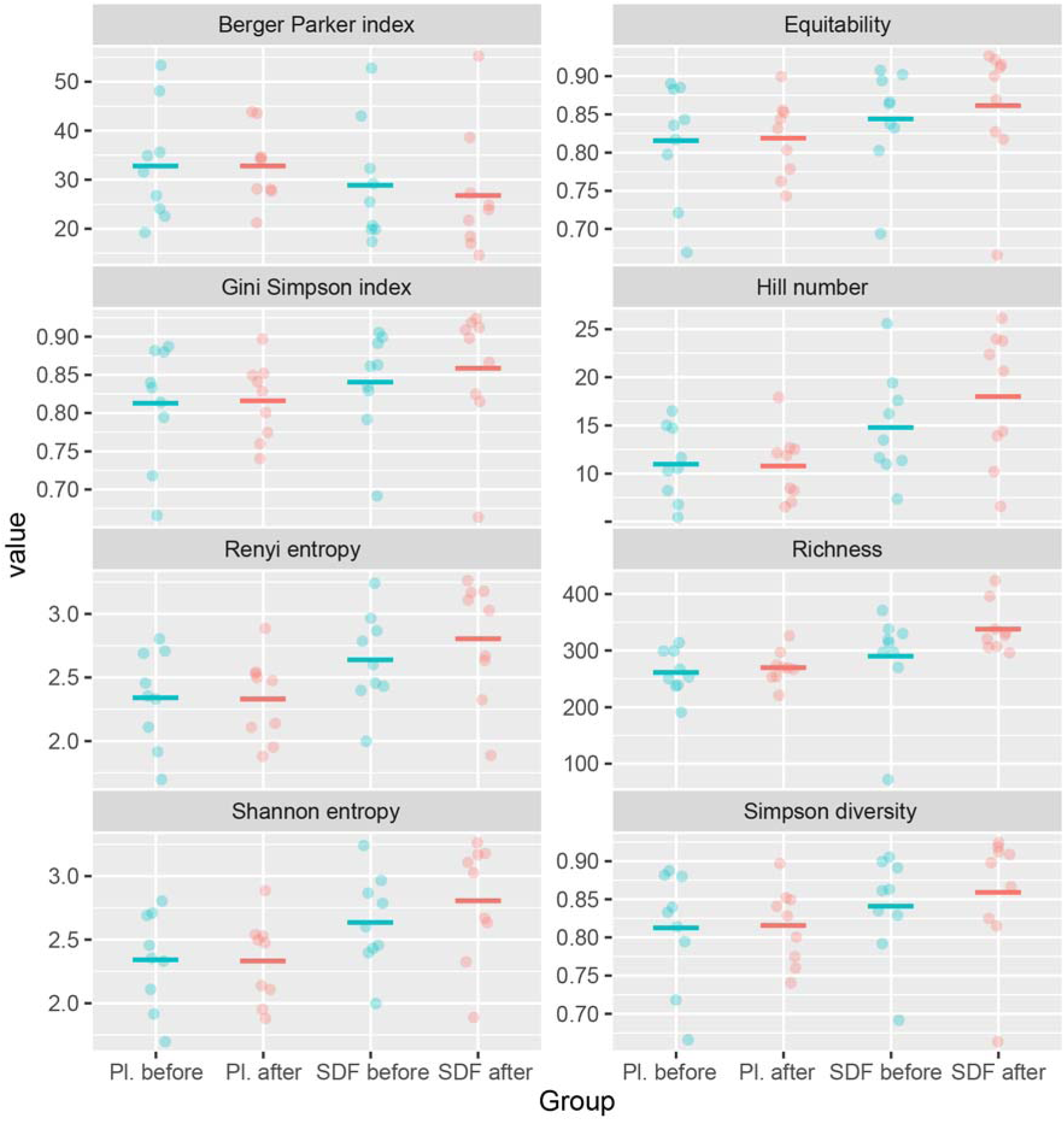
Various diversity analyses consistently assign no significant changes in microbial diversity following SDF treatment. A trend towards increased diversity is consistently observed. Pl: Placebo; SDF: Silver diamine fluoride.

**Figure S2.**
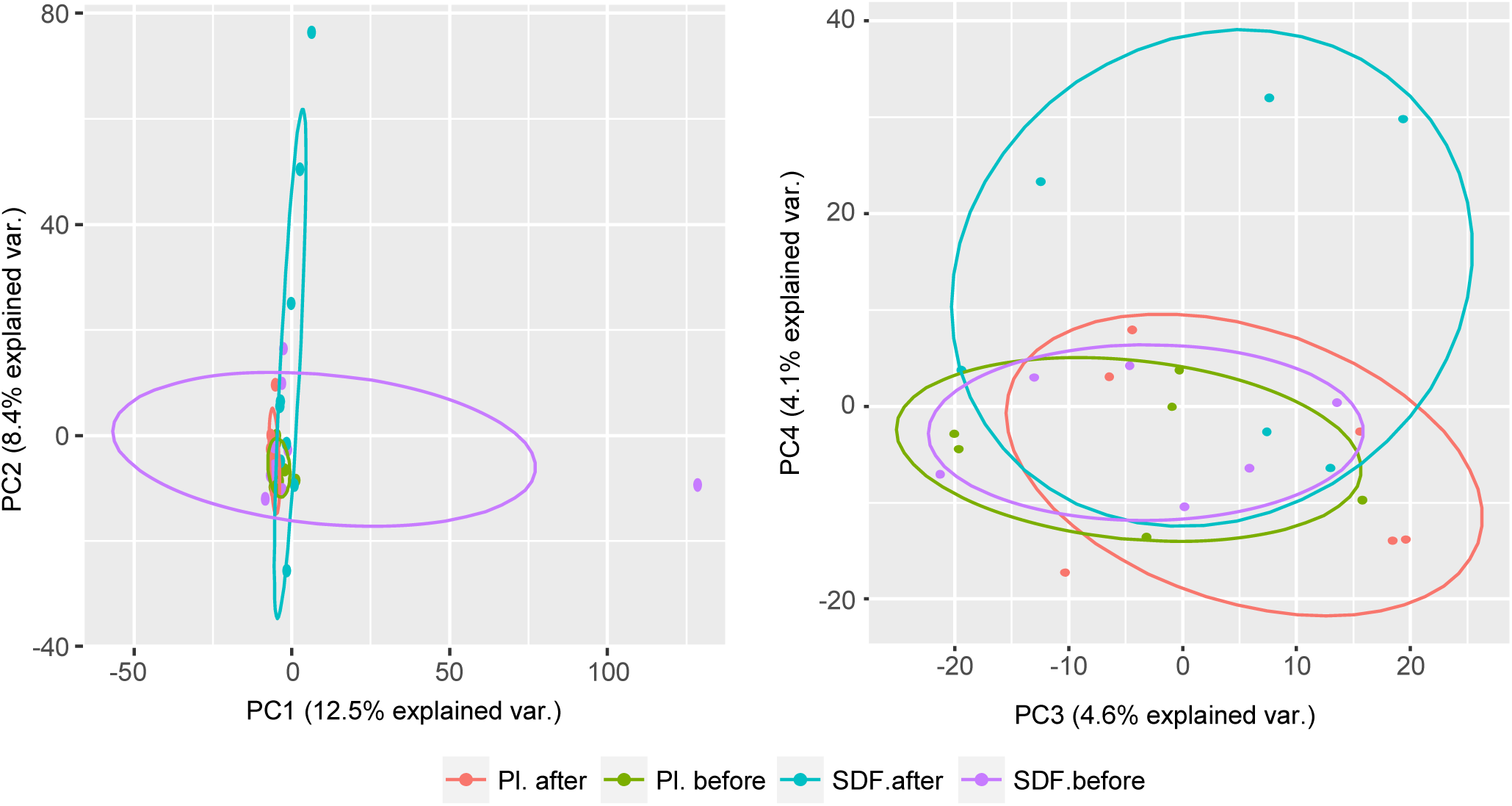
Principal component analysis shows outgrouping of half the post-treatment SDF group by components 2 and 4, which suggests subtle but distinguishable changes from treatment. The highly distributed explanation model (low attribution to any component) indicates high levels of similarity, or highly complex differences, between all samples. PC: principal component; Pl: Placebo; SDF: Silver diamine fluoride.

**Figure S3.**
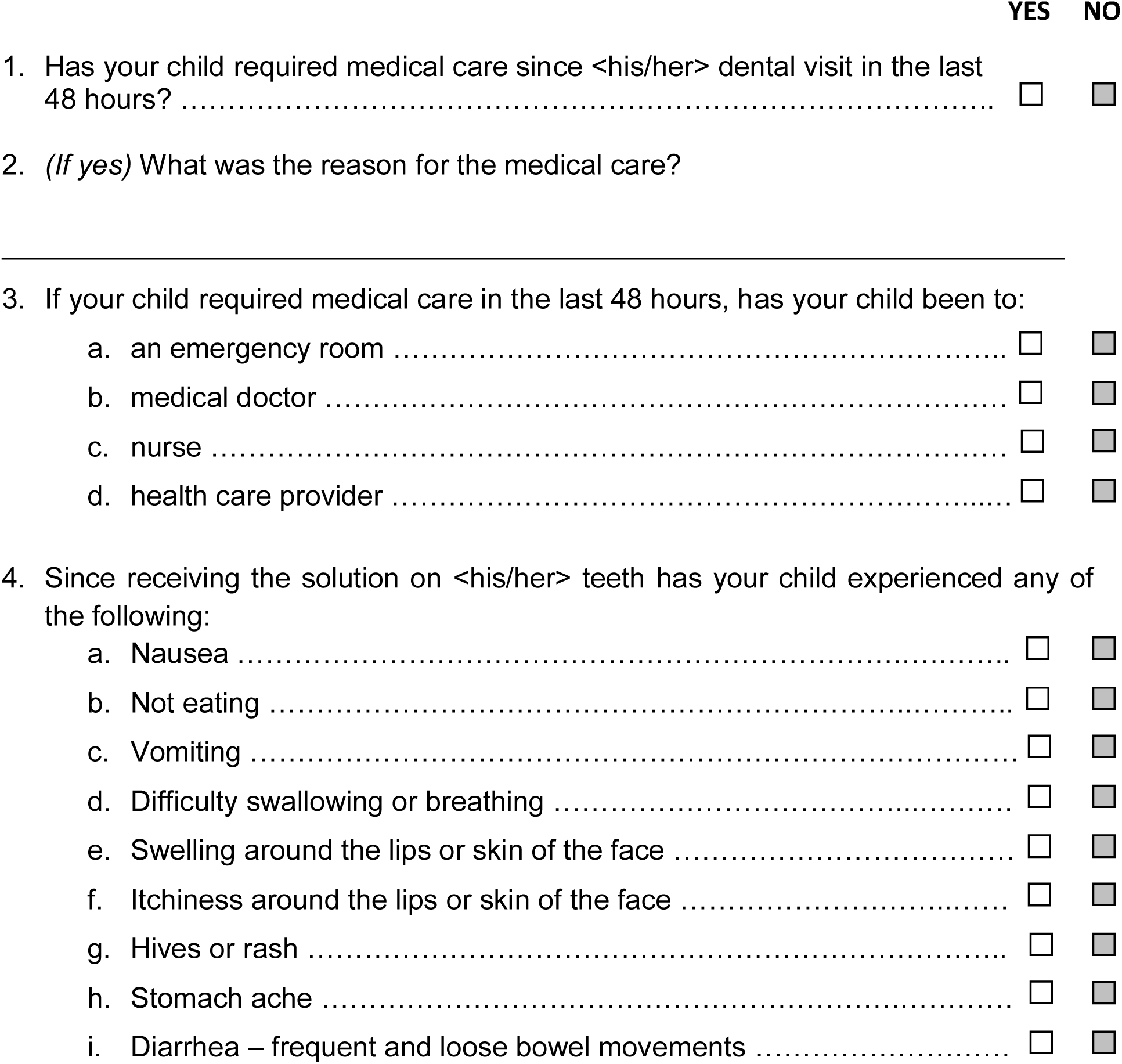
Safety questionnaire.

